# Visual Identification of Conspecifics Shapes Social Behavior in Mice

**DOI:** 10.1101/2024.06.07.597632

**Authors:** Devon Greer, Tianhao Lei, Anna Kryshtal, Zach Jessen, Gregory William Schwartz

## Abstract

Recognizing conspecifics in order to determine how to interact with them appropriately is a fundamental goal of animal sensory systems. It has undergone selective pressure in nearly all species. Mice have a large repertoire of social behaviors that are the subject of a rapidly growing field of study in neuroscience. Mouse social interactions likely incorporate all available sensory modalities, and the vast majority of studies have not attempted to isolate them. Specifically the role of vision in mouse social interactions remains unclear. We developed a behavioral platform that allowed us to present a subject mouse the visual information of stimulus mice in isolation from olfactory, acoustic, and tactile cues. Our results indicate that the visual identification of the sex or individual identity of other mice influences behavior. These findings highlight the underappreciated role of vision in mouse social interactions and open new avenues to study the visual circuits underlying social behavior.

## Introduction

For hundreds of millions of years before the evolution of the cerebral cortex, innate, hard-wired visual behaviors in animals contributed to their ability to survive core adaptive pressures: aligning activity and sleep to different parts of the circadian cycle^1,2^, finding food^3^, avoiding predators^4–6^, and interacting with conspecifics^7^. Humans have retained a remarkably diverse set of connections from the retina to dozens of subcortical targets in the brain^8–11^ – many associated with behaviors like hunger^12,13^, social-related fear^14–22^, mood^23,24^, and sexual attraction^25^ – yet we know almost nothing about the neural circuits in these pathways or their, presumably subconscious, influences on behavior.

The classic examples of specific retina-to-brain circuits influencing behavior come from “non-image forming” vision. Intrinsically photosensitive (ip)RGCs control circadian photoentrainment^26^, the pupillary light reflex^26^, and even mood^27^, each through a distinct circuit^28^. Much of the work connecting retinal cells to non-image forming behavior over the last decade was driven by papers introducing two new paradigms for studying innate visual behavior in mice: (1) escape responses to “looming” objects overhead^29^ and (2) predation of crickets^30^ or moving cricket-like dots^31^.

Along with avoidance of predators and identification of prey, recognition of conspecifics is one of the core visually guided behaviors under selective pressure. In studying predator avoidance and prey detection, there has been substantial progress in the last decade identifying the relevant stimuli and circuits involved^32–34^. While social behaviors are a central focus of daily life, in contrast, very little is known about the stimuli capable of driving visually guided social behaviors. We report evidence that vision plays a role in the recognition of conspecifics in several contexts.

Social recognition distinguishes familiar from unknown individuals of one’s own species. The ability to recognize a familiar individual is the foundation for social relationships, including parent-offspring recognition, mate recognition, and dominant-subordinate hierarchies. Comparatively little work has examined the role of vision in social behaviors in rodents, where olfaction has long been assumed to be the dominant sense^35^. There is, however, some dispersed evidence that vision plays a role in rodent social identification. Virgin male mice attack silicone objects only when they are shaped like pups, even in the absence of olfactory cues^36^. Male rats demonstrate non-contact penile tumescence in the presence of females at distances where odor cues were weak and did not demonstrate penile tumescence to receptive female odor alone^37^.

In the same way that looming and cricket-tracking experiments in mice led to major discoveries about the cell types and circuits involved in those behaviors^32–34^, our new behavioral paradigms will lay the foundation for work to reveal the circuits driving visual components of social behavior. Future studies building on our results could reveal how these circuits might be altered in social disorders like autism spectrum disorder.

## Results

### A platform for exploring visually-driven social behavior isolates vision from sound, smell, and touch

Our strategy to investigate visual cues driving social behavior was to isolate vision from other sensory modalities. We created a social behavior arena containing a central ring for the “subject” mouse and three isolated chambers with independent air supply to place stimulus objects, including other mice (**Figure 1**). We tracked the position and pose of the subject animal with four cameras: one above the arena and three embedded in the side walls. We recorded the mouse’s ultrasonic vocalizations with a microphone above the arena.

**Figure 1.**
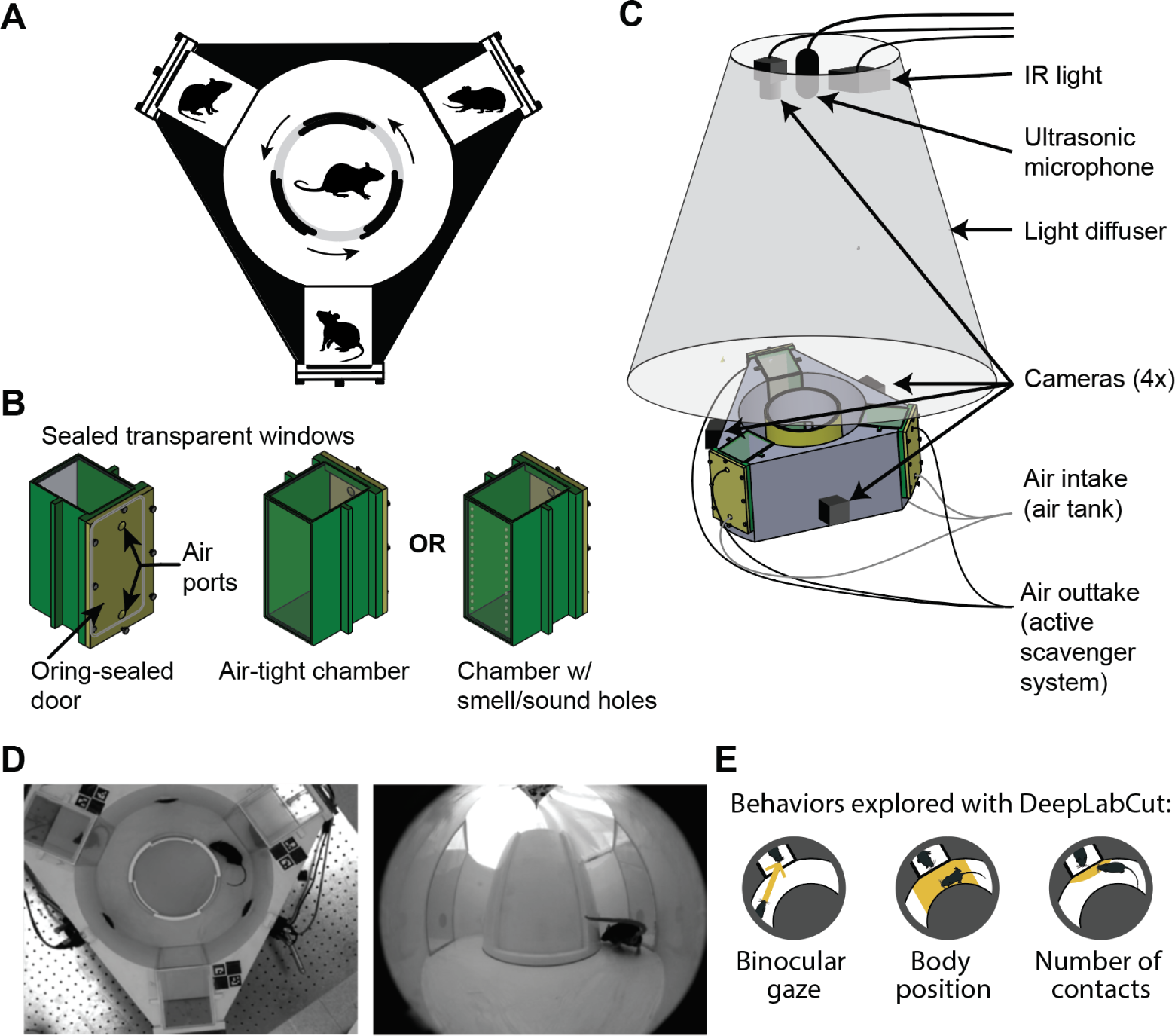
Behavior platform explores mouse visual social behavior by removing other social cues. **(A)** Schematic top view of the arena. **(B)** Stimulus chambers were O-ring sealed shut and slid into place within the platform. Air ports on the chamber doors connected to an air tank and an active scavenging machine. Stimulus chambers could be sealed to isolate smell and sound of the stimulus mouse or switched with others with perforations in control experiments. **(C)** Platform consisted of 4 cameras (1 top-down camera and 3 side view fisheye cameras embedded in the platform), air intake and outtake for the mice in the stimulus chambers, a light diffuser covering the top of the platform, IR light, and ultrasonic microphone. The inner annulus doors slid open to the outer annulus through a crank connected through the bottom. **(D)** Example frames captured simultaneously from two cameras. Left: top down camera. Right: Side camera with fish-eye lens. **(E)** Behaviors explored using DeepLabCut: binocular gaze (quantified by the number of seconds the observer mouse looked directly at the stimulus mouse), body position (determined by how many seconds the mouse was in front of a stimulus window), and by counting the number of contacts the mouse made onto the stimulus window (see **STAR Methods**).

We used DeepLabCut^38^ to track 12 locations on the animal, including the nose, ears, skull base, hips, feet, tail base, and tail tip (see **STAR Methods**). We used these measurements to compute three metrics: (1) direction of binocular gaze (2) body position, and (3) contact events when the nose touched one of the windows. In habituation sessions, the subject mice viewed empty chambers (see **STAR Methods**). None of the three response metrics showed preferences for one window over another in the habituation sessions (p = 0.5562, p = 0.4763, p = 0.4489, respectively).

To confirm that the chambers isolated sound from vision, we compared the recorded vocalizations between a condition in which a dam and her pups occupied the arena and a condition with the same number of pups present but isolated in the chambers. Pups produce quiet but frequent and predictable vocalizations with frequencies between 30 and 90 kHz^39–41^. The number of vocalizations was 6-10 fold higher when the pups were inside the main arena than when they were in the chambers (p = 0.0043, **Supplementary Figure 1**), demonstrating effective sound isolation. The calls recorded with the pups inside the chambers can be attributed to the dam in the main arena. Consistent with this interpretation, there was a higher proportion of “adult only” call types (see **STAR Methods**) and very few calls when the subject mouse was a male viewing pups (**Supplementary Figure 1)**.

Previous studies have shown that mice prefer conspecifics over novel objects^42–46^. To verify that mice were using visual information from the stimulus animals in the chamber rather than another cue, like odor, we placed a cagemate in one window and novel, inanimate objects in the other two windows (**Figure 2A**), and we compared normal light conditions to complete darkness. We tracked the subject mice from the top-down camera, and determined how much time they spent in front of the cagemate (**Figure 2B-C**) compared to the objects. The mice (male, n = 17) spent more time near the cagemate (body position, p = 0.0077) than the novel objects, and they faced the cagemate for more time (p = 0.0028) than towards the objects (**Figure 2D-F**). Subject mice made more contacts to the window containing their cagemate as opposed to the objects (p = 0.0312). In contrast, in control experiments in which 17 male mice in the arena were placed in complete darkness, all three effects were eliminated (p = 0.9482, 0.5379, 0.7663), indicating that they were driven by vision (**Figure 2G-I**).

**Figure 2.**
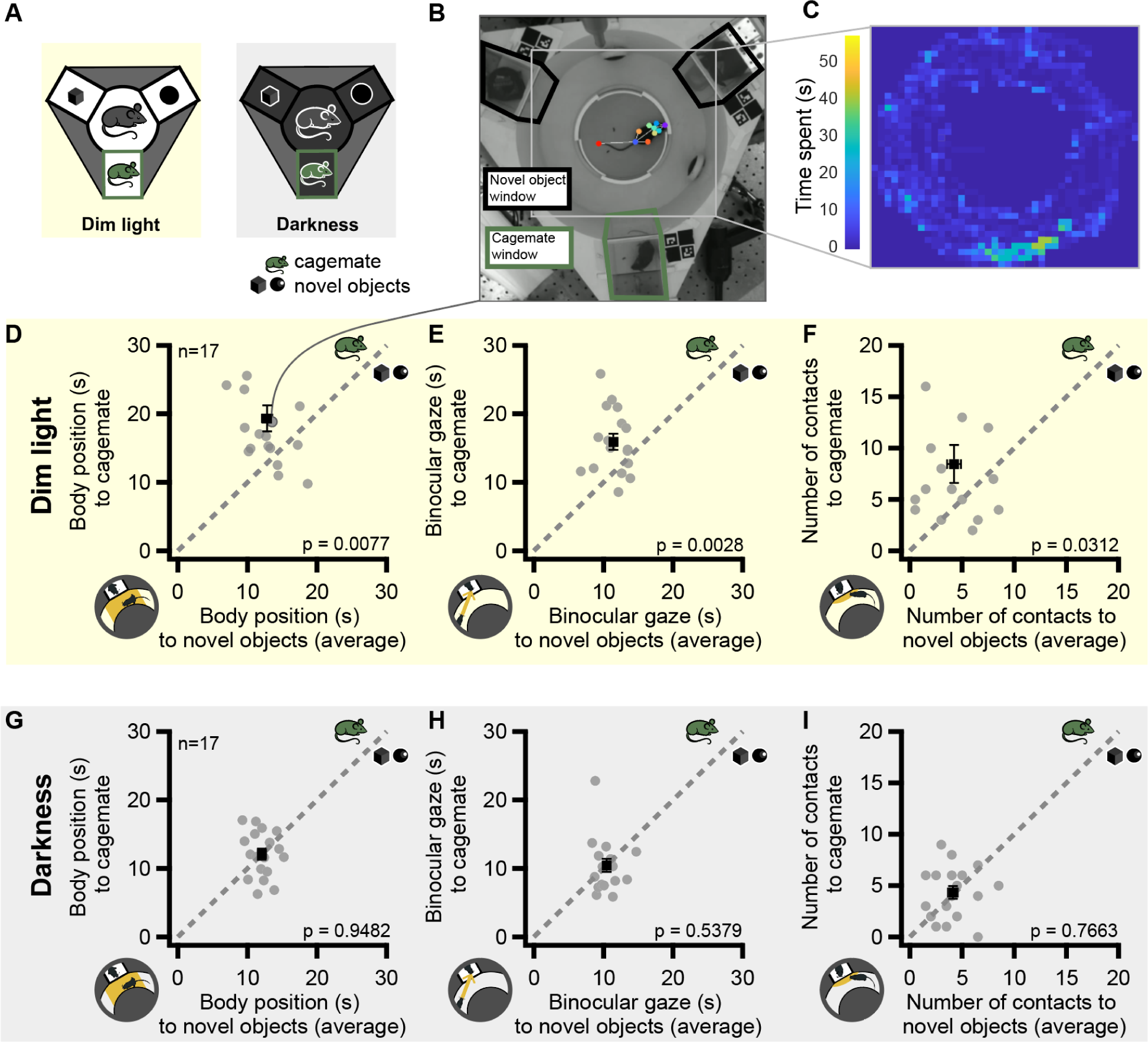
The platform isolates vision from sound, touch, and odor social cues. **(A)** Left: Experiment setup in ambient light conditions, mice viewed a cagemate (green) and two novel objects (black) in the 3 stimulus windows. Right: experimental conditions with a cagemate and two novel objects in complete darkness. **(B)** Example experiment in dim light conditions^47^, shown in dark gray outlines. The green border indicates the cagemate window and the black outlines the novel object locations. The light gray rectangle represents the area covered in **(C). (C)** Heatmap of the body position of the mouse in the example session. **(D)** The total number of seconds each mouse spent near the window with a cagemate compared to the seconds on average between the two novel object windows. Gray dashed lines indicate the unity line between the two stimulus types. P-values comparing this metric for the cagemate window vs. the other windows is indicated. Data point with a gray line extending to **(B)** indicates where the data from **(B)** falls on the scatter plot. **(E)** Comparing the number of seconds the mouse was directly facing their cagemate to the mean of the seconds spent facing the novel objects. **(F)** Number of contacts to the window of a cagemate as opposed to the two novel object windows. **(G-I)** Same as **(D-F)** but in complete darkness.

### Mice respond differently to the sight of male vs. female conspecifics

Having verified that the chamber effectively isolates vision from the other senses, next we tested whether the sex of the stimulus mice could influence the behavior of the subject mouse. We presented both male and female mice (24 males, 24 females) with two different ratios of males and females. The chambers either had one male and two females or one female and two males (**Figure 3A** and **Figure 3E**).

**Figure 3.**
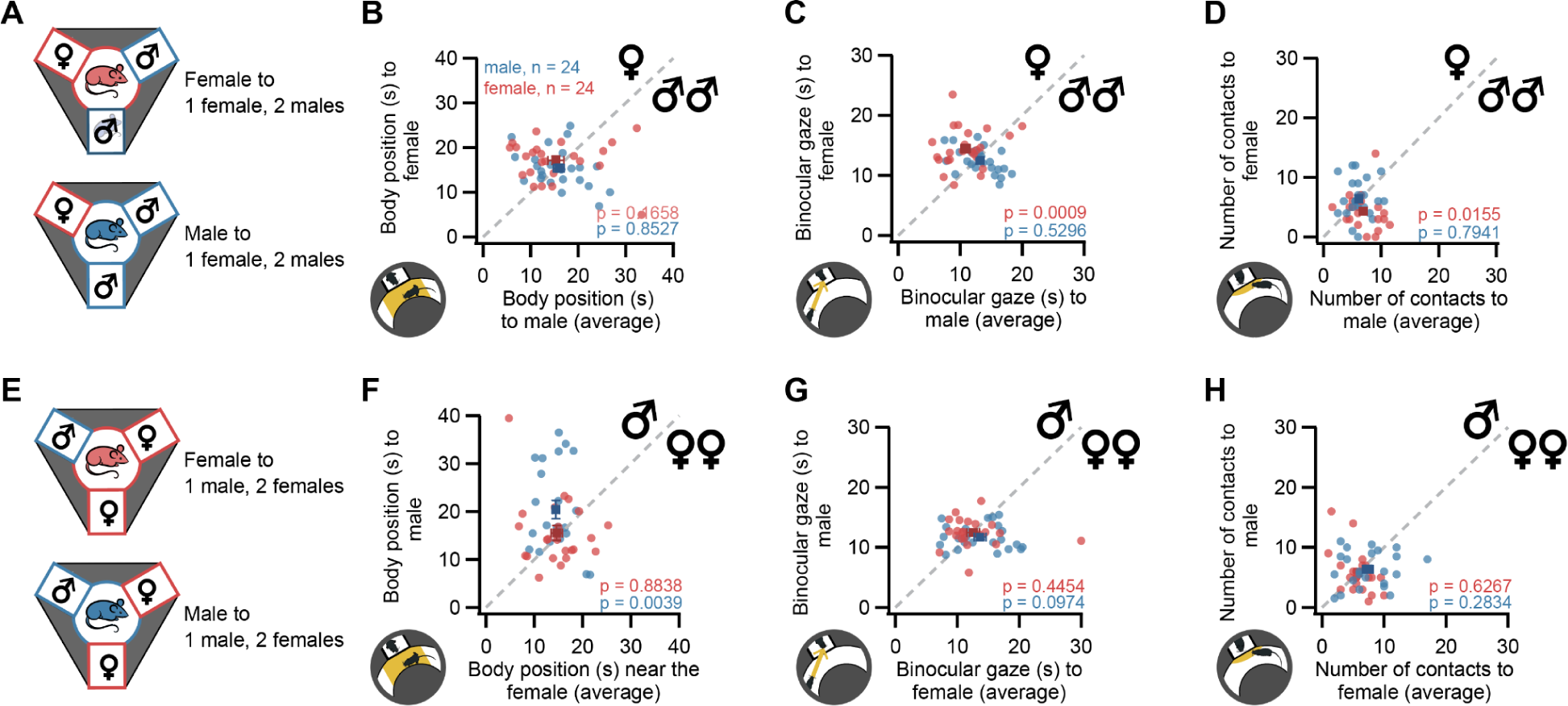
Mice recognize the sex of another mouse using vision alone. **(A)** Upper panel: female mice (red) and lower panel: male mice (blue) viewed **(B)** Body position time between the female versus the two male strangers (males n = 24, females n = 24). **(C)** Time facing the female compared to the two male strangers. **(D)** Number of contacts to the window with the female compared to the two males. **(E)** Experiment conditions in which mice viewed 1 female (red) and 2 males (blue) in the stimulus chambers. Upper panel: female mice (red) and lower panel: male mice (blue) viewed one female and two males in the platform windows. **(F-H)** The same metrics were explored as from **(B-D)** except with the experimental conditions being in front of the male stranger compared to the female strangers (averaged).

In conditions where one male and two females were in the stimulus chambers (**Figure 3A-D**), we found that females spent more time facing the male stranger compared to the female strangers (**Figure 3C**, p = 0.0009), while male observers exhibited no preference (p = 0.5296, using Wilcoxon signed-rank tests for comparison). We also found that the female observers made more contacts to the window of the male mice (p = 0.0155). In conditions with one male and two females (**Figure 3E-H**), we found that only the male subjects spent more time in proximity to the singular male mouse (**Figure 3F**, p = 0.0039), while the female observers showed no preference (p = 0.8838). These effects were similar when considering not just the central point of gaze, but either the binocular zone or the full field of view **(Supplementary Figure 2A-D**).

### The presence and sex of an intruder animal affects pup retrieval in dams

While these results demonstrated the ability for mice to recognize the sex of other mice using vision alone, next we wanted to explore whether this information has an impact on a strong, innate behavior. Maternal aggression and pup retrieval are highly stereotyped aggressive and defensive behaviors in female mother mice (dams) to protect their pups from intruders. Male intruders pose the greatest threat to the dam’s offspring^48^. The level of aggressive bouts is determined by the intruder’s sex^48,49^, with males receiving more attacks from the resident dam. In rats, the level of maternal aggression a male intruder receives is directly proportional to the intruder’s size^50^. To isolate the potential effects of competing interactions, we presented different-sexed intruder stimuli in separate behavioral sessions. We then examined whether the visual presentation of a male or female intruder would drive dams to retrieve their pups differently based on proximity to the intruder.

Each dam, their pups, and a hut from the home cage, were placed in the central area of the platform with the door open initially. A 30-minute acclimation period allowed the dam to groom, explore, and feed her pups (while the stimulus chambers remained empty). After this acclimation period, the dam was briefly enclosed in the central arena while either (1) a male stranger, (2) a female stranger, or (3) a novel object was placed in one of the stimulus chambers; the other chambers remaining empty (**Figure 4A**). The pups were then distributed evenly in front of the three windows in the outer arena. Recording commenced as the dam was released into the entire arena with access to her pups. We measured the order with which the dam retrieved pups in relation to the window behind which the intruder was situated.

**Figure 4.**
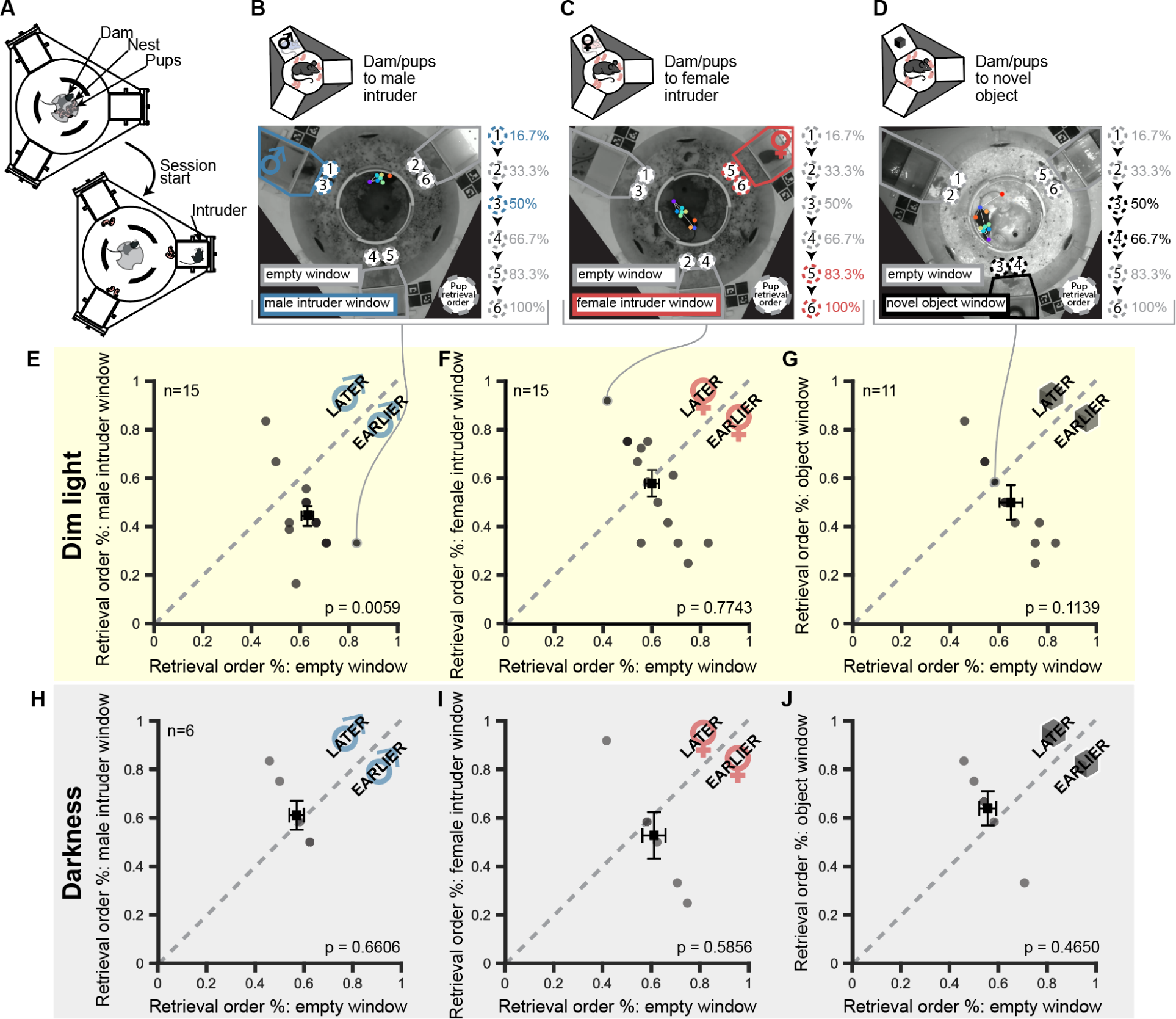
Sex of an intruder affects the maternal pup retrieval order. **(A)** Pup retrieval experimental procedure with a dam and pups is initiated first with acclimation to the platform with all viewing chambers empty. Pup retrieval was initiated distributing the pups evenly in front of the windows, with the arena divider closed, and the dam contained in the inner circle. An intruder was placed into one of the stimulus chambers. The arena divider was opened, allowing the windows to be visible to the dam. **(B)** Upper panel: experiment conditions for dam retrieving pups in the presence of a male intruder in one of the windows. Lower panel: blue outline marks the male intruder window in an example session and gray outline indicates empty windows. Circles with dashed outlines represent pups, with color representing if the pup is in front of a male intruder (blue) or empty (gray) window. Numbers 1-6 mark the order each pup was retrieved. Percentages indicate the percent of pup retrieval completed upon retrieval of that pup. **(C)** Experiment set up for pup retrieval in the presence of a female intruder. Same representations as **(B)** apply, except all intruders are female (red). **(D)** Experiment conditions for dam retrieving pups in the presence of a novel object in one of the windows. Same descriptions apply as with **(B,C)** except that novel object windows are indicated with black outlines. **(E)** In ambient light conditions, the order (by percent completed) that the pups in front of the male intruder window were retrieved compared the pups in front of the empty windows (mean) for n = 15 dams. Pup retrieval order was described as a percentage. **(F)** For sessions with female intruders, the same analysis was performed as **(E)** but for pup retrieval order in front of female intruders. **(G)** Sessions in front of novel object pup retrieval order compared to pup order in front of empty windows. Same results descriptions apply as for **(E, F)** however conditions were in the presence of a novel object. **(H-J)** For conditions, n = 6 dams, the same variables and metrics were explored as **(E-G)** however the experiment took place in complete darkness.

The sex of the intruder significantly influenced which pups the dams prioritized retrieving. When a male intruder was present, the dams exhibited a clear tendency to retrieve pups positioned closest to the male intruder earlier in the retrieval sequence compared to pups in front of empty windows (**Figure 4B**, p = 0.0059). In contrast, when a female intruder was present, there was no significant effect on pup retrieval order (**Figure 4C**, p = 0.7743). Similarly, the visual presentation of a novel object control had no significant impact on the dams’ retrieval order (**Figure 4D**, p = 0.1139). These findings demonstrate that the dams’ stereotyped maternal behavior of pup retrieval was modulated by the specific visual cue of a male intruder, prompting them to preferentially retrieve pups in closer proximity to this potential threat earlier in the sequence.

For comparison, we repeated these same experimental conditions, but in complete darkness. The dams (n = 6) appeared to not be influenced by the proximity of any intruder or unfamiliar object in one of the chambers. In these conditions, the same behavioral metrics explored produced insignificant effects from the presence of an intruder: body position (p = 0.6606), binocular gaze (p = 0.5856), or contacting the window (p = 0.4650).

### Mice recognize their cagemates compared to age and sex matched strangers

Previous literature has shown varied evidence regarding mice having a preference for novel mice over familiar ones, and this may depend upon context (and what is defined as a “familiar” mouse)^51–54,55^. However the literature is in agreement that mice are capable of making the distinction between familiar and unfamiliar conspecifics. We sought to investigate whether the identification of familiar mice could be driven using only visual information.

For these experiments, we measured the behavioral responses of mice (female n = 24, male n = 24) viewing their cagemate and two age and sex matched strangers within the chambers. The same three response metrics indicated that the mice could recognize their cagemates using vision alone. Males spent more time near the cagemate than near the strangers (**Figure 5B**, p = 0.009). Both males and females spent more time looking at the cagemate than the strangers (**Figure 5C**, p = 0.0081 in females, p = 0.0401 in males). Females made more contacts to the window of the cagemate than to that of the strangers (**Figure 5D**, p = 0.0160).

**Figure 5.**
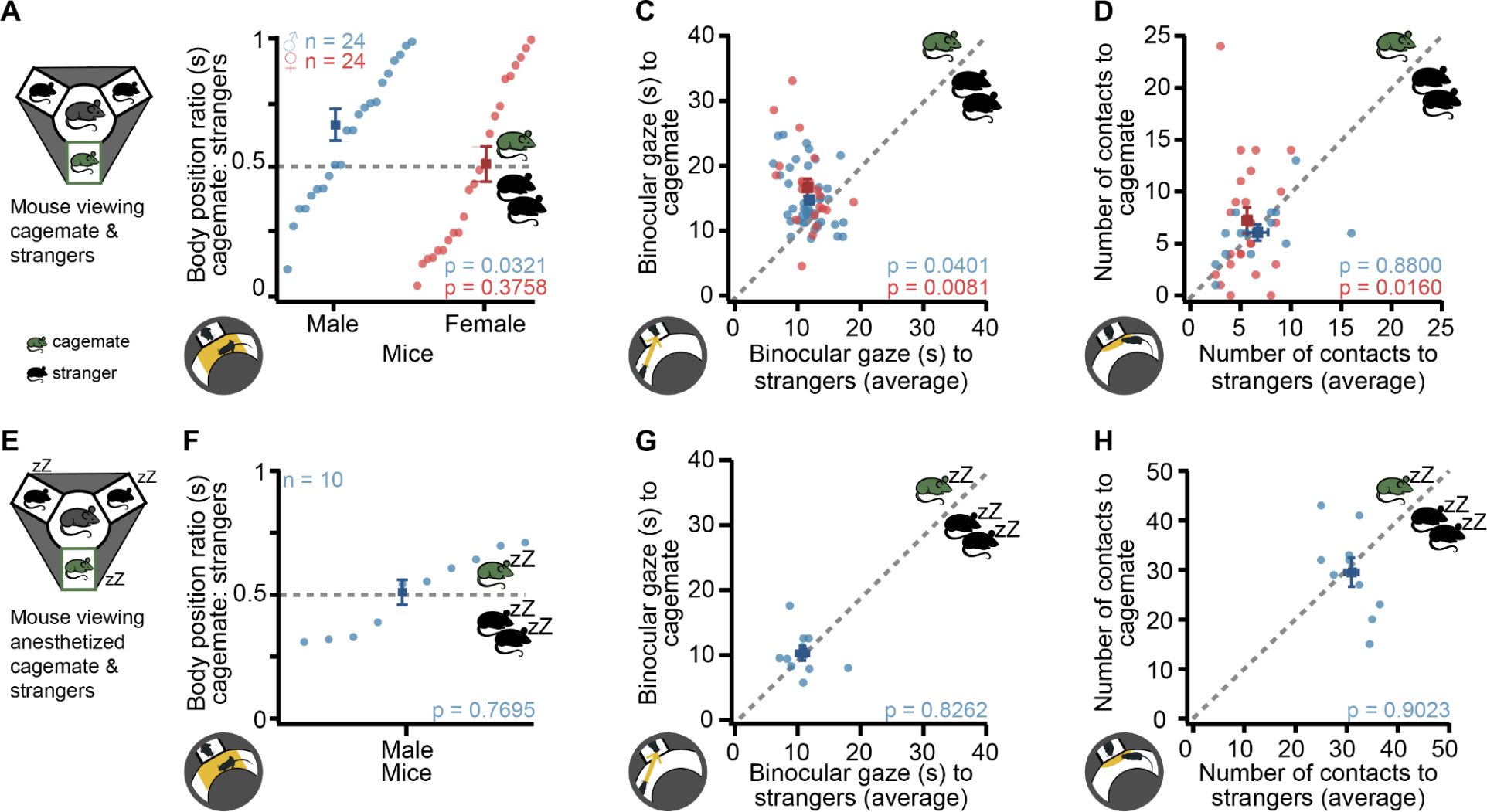
Familiar mice are identified from age and sex matched strangers. **(A)** Behavior set up in which mice viewed their cagemate and two sex and age matched strangers in the stimulus chambers. Green mouse icons indicate the cagemate, while black represent sex and age-matched strangers. **(B)** body position towards a cagemate compared to the strangers (average) for male and female mice (male n = 24, female n = 24). **(C)** Binocular gaze to their cagemate versus to the two strangers. **(D)** Number of times contact was made with the window with the cagemate compared to contacts with the windows with the strangers. **(E)** Same experimental organization applies as for **(A)**, except that all mice in visual chambers (cagemate and strangers) were anesthetized. **(F-H)** Same as **(B-D)** for behavior metrics but with the knocked out cagemate compared to knocked out strangers.

In accompanying experiments, we examined how the recognition of social cues of familiarity may depend on social status. Subordinate animals pay closer attention to the behaviors and cues of dominant group members. Attending these signals can be crucial for navigating the social hierarchy and avoiding conflict, as well as showing less aversion towards social cues from familiar subordinate individuals^56,57^. This attentiveness of lower-ranked individuals to dominant social signals has been well documented across a range of species^58–60^.

We tested both dominant and submissive mice with the visual cues of cagemates compared to strangers (**Supplementary Figures 3** and **4**) by manipulating the variables of familiarity (cagemates compared to strangers) and dominance status (dominant versus subordinate). We tested whether the observers’ behaviors were primarily driven by learned individual profiles of their cagemates or generalized visual signatures associated with contrasting dominance phenotypes. The time spent viewing the stimulus mice within their entire binocular field or visual field was also explored, **Supplementary Figures 5** and **6**. These results indicate they were able to extract and respond to generalized visual signals of dominance status, despite lacking direct experience with those specific individuals, however the submissive mice were more likely to recognize their familiar dominant counterparts.

### Motion cues are involved in cagemate identification

It is possible that the mice became attuned to their cagemates’ specific movement patterns through prolonged cohabitation. To decouple the potential influence of motion cues from other static visual signals, we repeated the experiment with the stimulus mice rendered immobile via sedation. By administering a ketamine/xylazine mixture to both cagemates and age/sex-matched strangers in the stimulus chambers, we eliminated any differential motion signals that could convey familiarity. Under these conditions where only static visual information was available, the subject mice no longer exhibited any preferences towards their cagemates compared to the stranger mice (**Figure 5F-H**).

## Discussion

### Isolating vision from other senses in freely behaving mice

In the realm of animal behavior, a critical objective of sensory systems is the identification of conspecifics to enable appropriate social interactions. However, the specific contributions of these modalities, particularly vision, have remained inadequately explored due to the integrative nature of previous studies. To address this gap, we engineered a behavioral platform specifically designed to isolate visual cues in the examination of social interactions among mice. This methodology permitted the presentation of visual stimuli from conspecifics devoid of olfactory, acoustic, and tactile information, thereby allowing us to dissect the visual component in social behavior.

From our controls with behavior in complete darkness (**Figure 2G-I & Figure 4H-J**), any preferences for engaging with the mouse stimuli over the novel objects were eliminated. We could interpret any effects found were due to visual cues associated with the presence of light. Experiments showed that mice use visual cues to identify the sex and individual identity of other mice. Mice behaved differently when presented with visual information from mice of varying sexes (**Figures 3** & **4**) and familiarity (**Figure 5**). These findings highlight the important but underappreciated role of vision in mouse social interactions and behavior. The results have implications for understanding social behaviors in mice and open new research avenues into the visual circuits mediating social behavior.

### Understanding the behavioral context and ethological relevance of visual-social behavior in rodents

Individuals keep alert to shifts in the social context of the group, such as alterations to group composition^59,61^. Across species, there is evidence that individuals respond differently to social cues based on factors like sex, hormonal state, past social experiences, and social status^62,63^. A mouse’s identity may influence how it perceives and responds to visual cues from conspecifics.

We found that mother mice prioritized retrieving pups (**Figure 4E**) located closest to a visually presented male intruder, which suggests they perceived the male as a potential threat to their offspring. By retrieving those pups first, the dams may have been motivated to remove the pups from the male’s vicinity. This behavioral adjustment based solely on visual cues demonstrates the mice’s ability to discern biologically relevant information through vision alone. Specifically, the male’s visual characteristics alone were sufficient to elicit altered maternal behavior.

Our findings underscore the importance of considering experimental context when assessing visually-driven social behavior in mice. The recognition of familiar mice appears to be context-dependent, with our results (**Figure 5)** suggesting that dynamic visual cues, such as motion signatures, may play a crucial role in individual recognition. The modulation of pup retrieval order by the visual presence of male intruders, but not females, demonstrates the ability of mice to extract biologically relevant information through vision alone and adapt their behavior accordingly.

Rapid assessment and anticipation^64^ of cagemate behavior patterns is crucial for survival and minimizing risks in the semi-naturalistic cage environment^65,66^. For instance, in mouse social groups containing an overtly aggressive individual, the ability to swiftly pick up on visual cues signaling that male’s temperament could be highly advantageous for subordinate cagemates. By closely monitoring dynamic visual signals, cagemates may be able to anticipate escalating aggression and adjust their own defensive responses proactively. This may grant a critical time advantage for mice to initiate avoidance or defensive actions before injurious conflicts erupt.

### Discerning the visual cues used to identify conspecifics

While the specific motivations behind these behavioral responses remain open to interpretation, the results demonstrate that mice are able to visually discern fundamental differences between male and female conspecifics (**Figure 3)** even in situations where no other sensory cues are available. The visual information was able to provide enough evidence that they could perceive sex-based characteristics through vision alone.

The visual features the mice could be using to extract social cues is worth exploring as stimulus design strongly shapes (and limits) neural responses^6,67^. While we do not expect them to be extracting cues with a high degree of acuity, there are a number of features they could use, including fur/skin colorations^68–70^, movement habits^71–73^, posture^74,75^, etc. The ability to discriminate familiars may have predominantly relied on learned visual motion signatures specific to each cagemate (**Figure 5)**. When dynamic cues were removed by sedating the stimulus mice, static visual features alone were insufficient for driving a divergent response between familiar and unfamiliar conspecifics in this context. This underscores how attuned the mice are to the minute kinematic patterns of their social partners through lifelong visual exposure and emphasizes vision’s role in encoding those individually distinctive motion profiles. Such continuous visual monitoring of familiars may result in a robust catalog of individualized motion and expression patterns.

As humans, we are also highly attuned to extracting social information from motion signatures^76^, and can also spot differences in sex^77^, emotion^78^, and identity^79–81^ based on motion signatures from a constellation of points alone. These features are unique enough to an individual to be used for biometric verification^82,83^. This highlights the importance of learned visual motion signatures in the social recognition process and this ability may be analogous to the way humans extract social information from motion cues.

### Brain circuits for processing visual social information

There are many reciprocal circuits by which visual social recognition could be being processed. We can narrow it down to the same pathways involved in social recognition, described as the triadic model^84,85^. These brain areas encompassing this process include the medial amygdala^85–87^, the prefrontal cortex^88^, and the striatum. The striatum’s role focuses on the motivation and decision-making processes^62^, while the medial prefrontal cortex contributes to the initiation, maintenance, and/or modulation of social behaviors^88^, and the medial amygdala (MeA) processes the emotionally and socially relevant information^89^.

MeA is known as a multisensory hub of social behavior^89–91^. Several papers have described a direct projection from the retina to MeA^92–97^. These retinal projections have been identified in mice^94,95,98,99^, rats^97,100^, hamsters^93,97^, cats^101^, and eight species of primates^102^. Several studies have suggested a visual component with links to MeA. Virgin male mice have characteristic aggressive behaviors toward pups that often result in the death of the infants^103^. This behavior relies on MeA, and it is present in the absence of olfactory cues (though touch was still available to the attacking males)^36^. Human studies have identified activation of MeA for faces making eye contact, even in a cortically blind individual^22^, and MeA lesion studies have shown impairments in the response to visual cues for sexual and emotional behaviors^104,105^.

### Developing new tools and paradigms for understanding social recognition

Recent advances in neuroscience have created powerful tools, computationally and genetically^106,107^, for studying neural circuits, revealing the complex mechanisms and pathways underlying social behavior^85,108–112^. Machine learning for body pose annotations can provide more opportunities for higher-level holistic, kinematic analysis of individual movements as a means of exploring new behavioral paradigms. The lack of novel behavioral paradigms is becoming a persistent barrier to making ethologically relevant findings^113,114^.

Our experiments demonstrate that mice can extract socially relevant information, such as sex and potential threat levels, through vision alone and modify their behaviors accordingly. They can distinguish mice based on sex (**Figures 3** and **4**) or familiarity (**Figure 5**) based solely on visual cues. This adds to a growing amount of literature showing that mice are capable of more with their vision for social cues. Their ability to process visual signals demonstrates how vision can profoundly shape motivated behaviors by providing basic information about the salience of social situations and stimuli in their environment. Our study provides evidence for the role of vision in social recognition and behavior in mice, highlighting the importance of considering experimental context and the dynamic nature of social interactions. These findings lay the groundwork for future research aimed at elucidating the neural circuits and mechanisms underlying visually-driven social behavior.

## Methods

The three cameras embedded within the walls of the platform had fisheye lenses to allow for full field-of-view coverage of the platform. The inner section of the platform was separated from the outer annulus of the platform by a sliding circular door, operated by a crank underneath the base of the platform. The walls between the inner and outer sections were at a slight incline to eliminate any obstruction of the viewing of the camera overhead. The three chambers allowed for the presentation of both objects or mice. They were supplied with an independent air source from a compressed air tank connected through air ports, along with a charcoal based active scavenging system. The platform was entirely covered by a one meter tall conical light diffuser. At the top of the light diffuser, the top-down camera, ultrasonic microphone, and infrared lights were mounted.

The cameras allowed us to monitor the position of the mouse at all times, from multiple angles. From these positions, we were able to estimate the body position of the mouse over the course of the session.

### Experimental model and subject details

#### Animals

Healthy adult 3–4 month old male and female C57BL6/J mice weighing 17-25 g (Jackson Laboratory, Bar Harbor, ME, USA) were housed in a 12 h normal light cycle for the duration of these studies. Food and water were available *ad libitum*, except for social dominance protocols described in the behavior methods section. After their arrival, mice were acclimated to the vivarium for a minimum of five days and then randomly assigned to experimental groups. All animal procedures were in accordance with relevant institutional and national guidelines and regulations and were approved by the Institutional Animal Care and Use Committee at Northwestern University.

#### Platform Specs

The test took place in an apparatus that was 3D printed at the university Research Core’s Design & Engineering Shop. Activity in the behavior platform was recorded via an overhead camera and three side cameras, Model: BFS-U3-13Y3M-C: 1.3 MP, 170 FPS, ON Semi PYTHON 1300, Mono (FLIR, Portland, OR, USA). The side cameras used fisheye lenses C-Mount 18 mm (Edmund Optics, Barrington, NJ, USA) to maximize visibility in the platform at the level of the mouse. Ultrasonic vocalizations were picked up on the Dodotronic Ultramic 250k ultrasonic microphone (Dodotronic, Italy). An air regulator distributed air into the stimulus chambers from a compressed air tank. The air outtake from the chambers was collected through an active scavenging system, R546-Pro (RWD, Sugarland, TX, USA). The arena was illuminated by 850nm infrared lights (Tendelux, China).

#### Habituation

Mice were brought into the testing room in their home cage and allowed to acclimate for at least one hour before any behavioral tests. Habituation took place in the behavior platform in low light conditions within 0-4 hours into the dark/active phase of their light cycle, respectively. Mice were placed in the arena and allowed to explore for five minutes^115^. After each session the arena was wiped with 70% ethanol and rinsed with water. Mice were never picked up by their tails during handling^116^.

#### Sex identification testing

After six days of habituation, mice were randomly assigned to experiment types. Those in the sex identification experiments each begin with the subject mouse being placed in the central arena of the platform, with the doors to the outer arena closed off. The stimulus mice, either 1) two females and one male or 2) two males and one female are placed in the three stimulus chambers. The subject mouse is released into the outer annulus of the arena using the crank underneath the platform. Once the mouse has entered the outer arena of the platform, the crank closes the central part of the arena from the outer once again. The subject mouse gets five minutes to explore the outer annulus of the arena, viewing the stimulus mice. Once five minutes is complete, the mice are removed from the platform.

#### Pup retrieval experiments

Prior to experiments, mice were assigned randomly to the pup retrieval experiments. Two female and one male mouse were placed in a cage for one week, then the male was removed from the cage. Female mice underwent habituation during gestation. Once the dams dropped their pups, the time/day was recorded. Upon postnatal age three to six days of the pups, the experiments were conducted. Bedding material from their cage was distributed throughout the platform. The dam, pups (three, six, or nine pups per dam), and the hut from their cage were placed in the central area of the platform, with the platform door closed. The crank was then turned to allow the mice access to all of the platform. The mice were given 30 minutes to become acquainted with the environment. At this point, the stimulus chambers were empty. The dam was monitored to confirm she was habituated sufficiently to groom, explore, and feed her pups during that time.

The experiment began with closing the mice into the central arena once again. The stimulus mice were then placed in the stimulus chambers. In this case, only one stimulus chamber receives one mouse, either 1) a male stranger, 2) female stranger, or 3) novel object control. The pups are then removed from the central arena and placed in equal numbers in front of each of the three windows in the outer arena. The recording was initiated when the dam was released into the arena from the center, allowing her access to her pups once again. The experiment was recorded for five minutes and then the mice were returned to their home cages. Outcomes measured included the order and latency with which the dam retrieved the pups in relation to the window the intruder was behind.

Experiments conducted in the dark had the same procedure but any light was extinguished once the pups were distributed in the arena prior to the recording. The platform doors were opened using the crank in total darkness. Using a verbal command on a phone app, a timer is commenced to keep track of the session duration. Once the countdown is reached, the computer monitor is turned back on to stop the recording.

#### Dominance experiments

To establish the necessary social contexts, we utilized male hierarchy pentads - small socially-organized groups of five males where a stable dominance hierarchy had naturally emerged over time, **Supplementary Figure 4**. From each nest group, we identified the most dominant and submissive individuals to serve as our experimental mice based on extensive behavioral observations of agonistic interactions, displacements, and other status-defining metrics. This was carried out in nine separate male hierarchies comprising five mice each.

When housed together, male mice form dominance hierarchies that can be assayed and manipulated with established methods^117^. To reinforce and solidify the dominant/subordinate phenotypes, we administered additional interventions over four consecutive days. For the dominant males, we provided exposure to novel, sexually receptive female mice, with receptivity confirmed through estrous cycle tracking. This positive reinforcement helps promote and maintain territorial, male-typical assertive behaviors^118,119^. Conversely, for the subordinate males, we implemented mild food restriction to bring their weight down to 90% of original levels and removed hair to represent barbering^117^.

To determine dominance hierarchies in mice, two types of methods are standardly used. In one, the dominance matrix is reorganized such that some numerical criterion, calculated for the matrix as a whole, is minimized or maximized. The other aims to provide a suitable measure of individual overall success, from which a rank order can be directly derived. Using a combination of both approaches, we monitored the highest and lowest members of already established groups, **Supplementary Figure 3**.

Sheets of precut Whatman grade 540 acid-hardened filter paper, WHA1540150, (Sigma-Aldrich, St. Louis, MO, USA) were cut to size and spread below each cage to collect urine deposited by the males. Male mice were maintained in these urine-marking cages for 4 h and then returned to their home cages. Urine marked filter paper was allowed to air-dry overnight and then the urine was visualized using ninhydrin (Sigma-Aldrich, St. Louis, MO, USA). Urine spots were stained with ninhydrin at a 2% dilution in 190-proof ethanol (Sigma-Aldrich, St. Louis, MO, USA) visualized with bright-field light. Dried Ninhydrin-stained sheets were scanned to a computer as TIF files. Urine void spots were identified by their purple coloration due to Ninhydrin staining and round shape; non-urine marks were identified as either fecal material (brown spots/smudges) or paw prints (purple paw-print shapes). These non-urine artifacts were erased in Adobe Photoshop (Adobe Systems, San Jose, CA, USA) prior to analysis. Urine patterns were quantified to determine which mice were more socially dominant. We counted both the number of individual urine events in both the outer perimeter of the cage and central 50% of the cage. Staining with ninhydrin allows clear visualization of the borders of overlapping voids however as a precaution, we additionally quantified the areas by pixels with urine markings. This was done in the Fiji version of ImageJ (http://fiji.sc/).

The Void Whizzard^120^ plugin from University of Wisconsin-Madison is an advanced tool for automated analysis of VSA images with several major advantages. It has been independently validated by several different laboratories. It is also able in most cases to automatically separate nonconcentric overlapping urine spots thus increasing data resolution and accuracy. Additionally, it is freely available as an ImageJ plugin with instruction manual (https://imagej.net/Void_Whizzard).

We counted (a) the total number of non-perimeter (interior) areas that contained any indication of urine marking, and (b) the number of central squares that contained any urine.

#### Cagemate identification testing

Following five days of habituation, mice were assigned randomly to the sex identification experiments. Each experiment commenced with the subject mouse being placed within the central arena of the platform, while the doors to the outer arena remained closed. The stimulus mice were placed in the stimulus chambers, with a cagemate in one chamber and two age and sex matched strangers in the other two chambers. The test mouse was introduced into the outer ring of the arena by operating the crank. Subsequently, after the mouse entered the outer arena, the crank sealed off the central section from the outer area once more. The test mouse was given five minutes to explore the outer ring of the arena and observe the stimulus mice. After the allotted five minutes have elapsed, the mice are taken out of the platform.

Experiments with anesthetized mice were conducted with the same procedure however prior to placement of the stimulus mice in the chambers, they were anesthetized with a intraperitoneal (IP) bolus of ketamine-xylazine (100 mg/kg and 10 mg/kg body mass, respectively) and confirmed to be knocked out testing for a toe pinch withdrawal. Breathing rate was monitored by eye at a rate of 2-3 Hz. Further 10%–20% of the ketamine-xylazine mixture was injected IP as necessary to maintain surgical plane anesthesia between sessions.

#### Statistical analysis and pose estimation

Statistical analysis was performed using Matlab and Python. Specific details for each experiment are listed in the associated figure legends. Wilcoxon signed-rank tests were performed to determine p values until otherwise described.

Pup retrieval behavioral interactions were scored manually using BORIS (Behavioral Observation Research Interactive Software) and reported as tabulated events. Pup order of retrieval was determined by the percentage of pup retrieval completion marked by returning any given pup.

Ultrasonic vocalizations (USVs) were detected using the ultrasonic microphone and using Deepsqueak machine learning. Classification was performed using supervised clustering and manual classifications. The .tdms files were converted to .wav format and imported into DeepSqueak for analysis. The following parameters were set for each file: total analysis length of 0, analysis chunk length of six, frame overlap of 0.0001 seconds, frequency low cut off of 30 kHz, frequency high cut off of 120 kHz, and score threshold of 0. To maximize the detection of ultrasonic vocalizations, the detection setting was set to “high recall.”

After the initial automated detection, the files underwent manual processing. This involved visually inspecting each detected USV to confirm it met previously established criteria for a true vocalization. Furthermore, the tonality threshold was manually adjusted for each file to optimize the signal-to-noise ratio. When necessary, the automatic detection boxes were redrawn to ensure they captured the full duration and frequency range of each vocalization.

The pose estimation of mice was conducted using the Python-based DeepLabCut toolbox, employing a deep neural network (NN) trained through transfer learning. We selected 10 key body parts for reconstructing the mouse skeleton: nose, left ear, right ear, neck base, left shoulder, right shoulder, left hip, right hip, tail base, and tail tip. Training of the NN was conducted using a dataset comprising 426 manually labeled 2D frames extracted from top-view camera videos, encompassing all experimental conditions within the behavior platform. The training process involved a total of 127,000 iterations. Notably, the training and test errors were measured at 2.42 pixels and 6.12 pixels, respectively. To enhance accuracy in labeling and eliminate outlier points, a cut-off threshold of 0.6 was employed.

## Data and code availability

- Behavior data have been deposited at Mendeley Data and are publicly available as of the date of publication. Accession numbers are listed in the key resources table.
- Schematic of the behavior platform has been deposited at Mendeley Data and is publicly available as of the date of publication. DOIs are listed in the key resources table.
- Any additional information required to reanalyze the data reported in this paper is available from the lead contact upon request

## Key Resources Table

| REAGENT or RESOURCE | SOURCE | IDENTIFIER |
| --- | --- | --- |
| Chemicals, Peptides, and Recombinant Proteins |  |  |
| Ninhydrin | Sigma-Aldrich | 151173 |
| 190-proof ethanol | Sigma-Aldrich | 493511 |
| Ketamine (80 mg/kg) |  |  |
| Xylazine (10 mg/kg) |  |  |
| Deposited Data |  |  |
| DLC Behavior data | Mendeley Data | DOI: 10.17632/br43h6z26t.1 |
| Three chambered mouse social behavior platform schematic | Mendeley Data | DOI: 10.17632/jxs94h9d5r.1 |
| Experimental Models: Organisms/Strains |  |  |
| model organism: C57BL/6J: B6 | Jackson Laboratory, ME, USA | Strain #:000664 |
| Software and Algorithms |  |  |
| Matlab | The MathWorks Inc., MA, USA | R2022b |
| ImageJ | NIH, USA <sup>121</sup> | https://github.com/fiji/fiji |
| DeepLabCut | Tübingen, Germany <sup>38</sup> | https://github.com/DeepLabCut/DeepLabCut |
| DeepSqueak | WA, USA <sup>122</sup> | https://github.com/DrCoffey/DeepSqueak |
| GitHub | Microsoft, WA, USA | https://github.com/SchwartzNU/DJ_schwartzlab |
| Adobe Photoshop | Adobe Systems, CA, USA |  |
| Void Whizzard | ImageJ plugin <sup>120</sup> | https://imagej.net/Void_Whizzard |
| Behavioral Observation Research Interactive Software | Torino, Italy <sup>123</sup> | https://github.com/olivierfriard/BORIS |
| Other |  |  |
| AI4 IR Illuminator, 4W Infrared Light | Tendelux | TX-AI4 |
| Blackfly S USB3 | FLIR, OR, USA | #BFS-U3-13Y3M-C |
| C-Mount 18mm fisheye lens | Edmund Optics, NJ, USA | #62-274 |
| Ultramic 250k ultrasonic microphone | Dodotronic, Gandolfo, Italy | SKU: UM250K |
| Whatman grade 540 acid-hardened filter paper | Sigma-Aldrich, MA, USA | WHA1540150 |

## References

1. LeGates, T.A., Altimus, C.M., Wang, H., Lee, H.-K., Yang, S., Zhao, H., Kirkwood, A., Weber, E.T., and Hattar, S. (2012). Aberrant light directly impairs mood and learning through melanopsin-expressing neurons. Nature 491, 594–598.

2. Tam, S.K.E., Hasan, S., Hughes, S., Hankins, M.W., Foster, R.G., Bannerman, D.M., and Peirson, S.N. (2016). Modulation of recognition memory performance by light requires both melanopsin and classical photoreceptors. Proc. Biol. Sci. 283. 10.1098/rspb.2016.2275.

3. Fleisch, V.C., and Neuhauss, S.C.F. (2006). Visual behavior in zebrafish. Zebrafish 3, 191–201.

4. Wei, P., Liu, N., Zhang, Z., Liu, X., Tang, Y., He, X., Wu, B., Zhou, Z., Liu, Y., Li, J., et al. (2015). Corrigendum: Processing of visually evoked innate fear by a non-canonical thalamic pathway. Nat. Commun. 6, 8228.

5. Evans, D.A., Stempel, A.V., Vale, R., Ruehle, S., Lefler, Y., and Branco, T. (2018). A synaptic threshold mechanism for computing escape decisions. Nature 558, 590–594.

6. Yilmaz, M., and Meister, M. (2013). Rapid innate defensive responses of mice to looming visual stimuli. Curr. Biol. 23, 2011–2015.

7. Dreosti, E., Lopes, G., Kampff, A.R., and Wilson, S.W. (2015). Development of social behavior in young zebrafish. Front. Neural Circuits 9, 39.

8. Hannibal, J., Hindersson, P., Ostergaard, J., Georg, B., Heegaard, S., Larsen, P.J., and Fahrenkrug, J. (2004). Melanopsin is expressed in PACAP-containing retinal ganglion cells of the human retinohypothalamic tract. Invest. Ophthalmol. Vis. Sci. 45, 4202–4209.

9. Fredericks, C.A., Giolli, R.A., Blanks, R.H., and Sadun, A.A. (1988). The human accessory optic system. Brain Res. 454, 116–122.

10. Hannibal, J., Kankipati, L., Strang, C.E., Peterson, B.B., Dacey, D., and Gamlin, P.D. (2014). Central projections of intrinsically photosensitive retinal ganglion cells in the macaque monkey. J. Comp. Neurol. 522, 2231–2248.

11. Sadun, A.A., Johnson, B.M., and Schaechter, J. (1986). Neuroanatomy of the human visual system: Part III Three retinal projections to the hypothalamus. Neuroophthalmology 6, 371–379.

12. Frank, S., Laharnar, N., Kullmann, S., Veit, R., Canova, C., Hegner, Y.L., Fritsche, A., and Preissl, H. (2010). Processing of food pictures: influence of hunger, gender and calorie content. Brain Res. 1350, 159–166.

13. Mora, F., Rolls, E.T., and Burton, M.J. (1976). Modulation during learning of the responses of neurons in the lateral hypothalamus to the sight of food. Exp. Neurol. 53, 508–519.

14. Pegna, A.J., Khateb, A., Lazeyras, F., and Seghier, M.L. (2005). Discriminating emotional faces without primary visual cortices involves the right amygdala. Nat. Neurosci. 8, 24–25.

15. de Gelder, B., Vroomen, J., Pourtois, G., and Weiskrantz, L. (1999). Non-conscious recognition of affect in the absence of striate cortex. Neuroreport 10, 3759–3763.

16. Hamm, A.O., Weike, A.I., Schupp, H.T., Treig, T., Dressel, A., and Kessler, C. (2003). Affective blindsight: intact fear conditioning to a visual cue in a cortically blind patient. Brain 126, 267–275.

17. Morris, J.S., Ohman, A., and Dolan, R.J. (1998). Conscious and unconscious emotional learning in the human amygdala. Nature 393, 467–470.

18. Morris, J.S., Öhman, A., and Dolan, R.J. (1999). A subcortical pathway to the right amygdala mediating “unseen” fear. Proceedings of the National Academy of Sciences 96, 1680–1685.

19. Breiter, H.C., Etcoff, N.L., Whalen, P.J., Kennedy, W.A., Rauch, S.L., Buckner, R.L., Strauss, M.M., Hyman, S.E., and Rosen, B.R. (1996). Response and habituation of the human amygdala during visual processing of facial expression. Neuron 17, 875–887.

20. Wright, C.I., Martis, B., Shin, L.M., Fischer, H., and Rauch, S.L. (2002). Enhanced amygdala responses to emotional versus neutral schematic facial expressions. Neuroreport 13, 785–790.

21. Zald, D.H. (2003). The human amygdala and the emotional evaluation of sensory stimuli. Brain Res. Brain Res. Rev. 41, 88–123.

22. Burra, N., Hervais-Adelman, A., Kerzel, D., Tamietto, M., de Gelder, B., and Pegna, A.J. (2013). Amygdala activation for eye contact despite complete cortical blindness. J. Neurosci. 33, 10483–10489.

23. Hannibal, J. (2006). Roles of PACAP-Containing Retinal Ganglion Cells in Circadian Timing. In International Review of Cytology (Academic Press), pp. 1–39.

24. Czeisler, C.A., Shanahan, T.L., Klerman, E.B., Martens, H., Brotman, D.J., Emens, J.S., Klein, T., and Rizzo, J.F., 3rd (1995). Suppression of melatonin secretion in some blind patients by exposure to bright light. N. Engl. J. Med. 332, 6–11.

25. Hamann, S., Herman, R.A., Nolan, C.L., and Wallen, K. (2004). Men and women differ in amygdala response to visual sexual stimuli. Nat. Neurosci. 7, 411–416.

26. Chen, S.-K., Badea, T.C., and Hattar, S. (2011). Photoentrainment and pupillary light reflex are mediated by distinct populations of ipRGCs. Nature 476, 92–95.

27. Fernandez, D.C., Fogerson, P.M., Lazzerini Ospri, L., Thomsen, M.B., Layne, R.M., Severin, D., Zhan, J., Singer, J.H., Kirkwood, A., Zhao, H., et al. (2018). Light Affects Mood and Learning through Distinct Retina-Brain Pathways. Cell 175, 71–84.e18.

28. Aranda, M.L., and Schmidt, T.M. (2021). Diversity of intrinsically photosensitive retinal ganglion cells: circuits and functions. Cell. Mol. Life Sci. 78, 889–907.

29. Yilmaz, M., and Meister, M. (2013). Rapid Innate Defensive Responses of Mice to Looming Visual Stimuli. Curr. Biol. 23, 2011–2015.

30. Hoy, J.L., Yavorska, I., Wehr, M., and Niell, C.M. (2016). Vision Drives Accurate Approach Behavior during Prey Capture in Laboratory Mice. Curr. Biol. 26, 3046–3052.

31. Allen, K., Gonzalez-Olvera, R., Kumar, M., Feng, T., Pieraut, S., and Hoy, J.L. (2022). A binocular perception deficit characterizes prey pursuit in developing mice. iScience 25, 105368.

32. Hoy, J.L., Bishop, H.I., and Niell, C.M. (2019). Defined Cell Types in Superior Colliculus Make Distinct Contributions to Prey Capture Behavior in the Mouse. Curr. Biol. 29, 4130–4138.e5.

33. Johnson, K.P., Fitzpatrick, M.J., Zhao, L., Wang, B., McCracken, S., Williams, P.R., and Kerschensteiner, D. (2021). Cell-type-specific binocular vision guides predation in mice. Neuron 109, 1527–1539.e4.

34. Wang, F., Li, E., De, L., Wu, Q., and Zhang, Y. (2021). OFF-transient alpha RGCs mediate looming triggered innate defensive response. Curr. Biol. 31, 2263–2273.e3.

35. de la Zerda, S.H., Netser, S., Magalnik, H., Briller, M., Marzan, D., Glatt, S., Abergel, Y., and Wagner, S. (2022). Social recognition in laboratory mice requires integration of behaviorally-induced somatosensory, auditory and olfactory cues. Psychoneuroendocrinology 143, 105859.

36. Isogai, Y., Wu, Z., Love, M.I., Ahn, M.H.-Y., Bambah-Mukku, D., Hua, V., Farrell, K., and Dulac, C. (2018). Multisensory Logic of Infant-Directed Aggression by Males. Cell 175, 1827–1841.e17.

37. Sachs, B.D., Akasofu, K., Citron, J.H., Daniels, S.B., and Natoli, J.H. (1994). Noncontact stimulation from estrous females evokes penile erection in rats. Physiol. Behav. 55, 1073–1079.

38. Mathis, A., Mamidanna, P., Cury, K.M., Abe, T., Murthy, V.N., Mathis, M.W., and Bethge, M. (2018). DeepLabCut: markerless pose estimation of user-defined body parts with deep learning. Nat. Neurosci. 21, 1281–1289.

39. Grimsley, J.M.S., Monaghan, J.J.M., and Wenstrup, J.J. (2011). Development of social vocalizations in mice. PLoS One 6, e17460.

40. Ehret, G. (1976). Development of absolute auditory thresholds in the house mouse (Mus musculus). J. Am. Audiol. Soc. 1, 179–184.

41. Branchi, I., Santucci, D., and Alleva, E. (2006). Analysis of ultrasonic vocalizations emitted by infant rodents. Curr. Protoc. Toxicol. Chapter 13, Unit13.12.

42. Tilly, S.-L.C., Dallaire, J., and Mason, G.J. (2010). Middle-aged mice with enrichment-resistant stereotypic behaviour show reduced motivation for enrichment. Anim. Behav. 80, 363–373.

43. Williams, C.M., Riddell, P.M., and Scott, L.A. (2008). Comparison of preferences for object properties in the rat using paired- and free-choice paradigms. Appl. Anim. Behav. Sci. 112, 146–157.

44. Garner, J.P. (1999). The aetiology of stereotypy in caged animals.

45. Mason, G.J., and Latham, N.R. (2004). Can’t stop, won’t stop: is stereotypy a reliable animal welfare indicator? Anim. Welf. 13, S57–S69.

46. Stevenson, M.F. (1983). The captive environment: its effect on exploratory and related behavioural responses in wild animals. Exploration in animals and man.

47. Richetto, J., Polesel, M., and Weber-Stadlbauer, U. (2019). Effects of light and dark phase testing on the investigation of behavioural paradigms in mice: Relevance for behavioural neuroscience. Pharmacol. Biochem. Behav. 178, 19–29.

48. Haney, M., Debold, J.F., and Miczek, K.A. (1989). Maternal aggression in mice and rats towards male and female conspecifics. Aggress. Behav. 15, 443–453.

49. Gammie, S.C. (2005). Current models and future directions for understanding the neural circuitries of maternal behaviors in rodents. Behav. Cogn. Neurosci. Rev. 4, 119–135.

50. Flannelly, K.J., Flannelly, L., and Lore, R. (1986). Post partum aggression against intruding male conspecifics in sprague-dawley rats. Behav. Processes 13, 279–286.

51. Moy, S.S., Nadler, J.J., Perez, A., Barbaro, R.P., Johns, J.M., Magnuson, T.R., Piven, J., and Crawley, J.N. (2004). Sociability and preference for social novelty in five inbred strains: an approach to assess autistic-like behavior in mice. Genes Brain Behav. 3, 287–302.

52. Wu, R., Jiang, X., Wu, X., Pang, J., Tang, Y., Ren, Z., Yang, F., Yang, S., and Wei, W. (2022). Interspecific differences in sociability, social novelty preference, anxiety- and depression-like behaviors between Brandt’s voles and C57BL/6J mice. Behav. Processes 197, 104624.

53. Hackenberg, T.D., Vanderhooft, L., Huang, J., Wagar, M., Alexander, J., and Tan, L. (2021). Social preference in rats. J. Exp. Anal. Behav. 115, 634–649.

54. Beery, A.K., and Shambaugh, K.L. (2021). Comparative Assessment of Familiarity/Novelty Preferences in Rodents. Front. Behav. Neurosci. 15, 648830.

55. Kareem, A.M., and Barnard, C.J. (1982). The importance of kinship and familiarity in social interactions between mice. Anim. Behav. 30, 594–601.

56. Borelli, K.G., Blanchard, D.C., Javier, L.K., Defensor, E.B., Brandão, M.L., and Blanchard, R.J. (2009). Neural correlates of scent marking behavior in C57BL/6J mice: detection and recognition of a social stimulus. Neuroscience 162, 914–923.

57. Lüscher Dias, T., Fernandes Golino, H., Oliveira, V.E.M. de, Dutra Moraes, M.F., and Schenatto Pereira, G. (2016). c-Fos expression predicts long-term social memory retrieval in mice. Behav. Brain Res. 313, 260–271.

58. Kunkel, T., and Wang, H. (2018). Socially dominant mice in C57BL6 background show increased social motivation. Behav. Brain Res. 336, 173–176.

59. Curley, J.P. (2016). Temporal pairwise-correlation analysis provides empirical support for attention hierarchies in mice. Biol. Lett. 12. 10.1098/rsbl.2016.0192.

60. Kingsbury, L., Huang, S., Wang, J., Gu, K., Golshani, P., Wu, Y.E., and Hong, W. (2019). Correlated Neural Activity and Encoding of Behavior across Brains of Socially Interacting Animals. Cell 178, 429–446.e16.

61. Williamson, C.M., Romeo, R.D., and Curley, J.P. (2017). Dynamic changes in social dominance and mPOA GnRH expression in male mice following social opportunity. Horm. Behav. 87, 80–88.

62. Chen, P., and Hong, W. (2018). Neural Circuit Mechanisms of Social Behavior. Neuron 98, 16–30.

63. Nikonov, A.A., and Maruska, K.P. (2019). Male dominance status regulates odor-evoked processing in the forebrain of a cichlid fish. Sci. Rep. 9, 5083.

64. Eilam, D., Izhar, R., and Mort, J. (2011). Threat detection: behavioral practices in animals and humans. Neurosci. Biobehav. Rev. 35, 999–1006.

65. Kavaliers, M., Colwell, D.D., Cloutier, C.J., Ossenkopp, K.-P., and Choleris, E. (2014). Pathogen threat and unfamiliar males rapidly bias the social responses of female mice. Anim. Behav. 97, 105–111.

66. Ginsburg, B., and Allee, W.C. (1942). Some Effects of Conditioning on Social Dominance and Subordination in Inbred Strains of Mice. Physiol. Zool. 15, 485–506.

67. Katsov, A.Y., and Clandinin, T.R. (2008). Motion processing streams in Drosophila are behaviorally specialized. Neuron 59, 322–335.

68. Lynn Lamoreux, M., Delmas, V., Larue, L., and Bennett, D. (2010). The Colors of Mice: A Model Genetic Network (John Wiley & Sons).

69. Jackson, I.J. (1994). Molecular and developmental genetics of mouse coat color. Annu. Rev. Genet. 28, 189–217.

70. Sundberg, J.P., and Silva, K.A. (2012). What Color Is the Skin of a Mouse? Vet. Pathol. 49, 142–145.

71. Bortolato, M., and Pittenger, C. (2017). Modeling tics in rodents: Conceptual challenges and paths forward. J. Neurosci. Methods 292, 12–19.

72. Tohda, C., Nakanishi, R., and Kadowaki, M. (2009). Hyperactivity, memory deficit and anxiety-related behaviors in mice lacking the p85alpha subunit of phosphoinositide-3 kinase. Brain Dev. 31, 69–74.

73. Langen, M., Kas, M.J.H., Staal, W.G., van Engeland, H., and Durston, S. (2011). The neurobiology of repetitive behavior: of mice…. Neurosci. Biobehav. Rev. 35, 345–355.

74. Miczek, K.A., Maxson, S.C., Fish, E.W., and Faccidomo, S. (2001). Aggressive behavioral phenotypes in mice. Behav. Brain Res. 125, 167–181.

75. Berman, G.J. (2018). Measuring behavior across scales. BMC Biol. 16, 23.

76. Blake, R., and Shiffrar, M. (2007). Perception of Human Motion (SSRN).

77. Kozlowski, L.T., and Cutting, J.E. (1978). Recognizing the gender of walkers from point-lights mounted on ankles: Some second thoughts. Percept. Psychophys. 23, 459.

78. Atkinson, A.P., Dittrich, W.H., Gemmell, A.J., and Young, A.W. (2004). Emotion perception from dynamic and static body expressions in point-light and full-light displays. Perception 33, 717–746.

79. Loula, F., Prasad, S., Harber, K., and Shiffrar, M. (2005). Recognizing people from their movement. J. Exp. Psychol. Hum. Percept. Perform. 31, 210–220.

80. Jokisch, D., Daum, I., and Troje, N.F. (2006). Self recognition versus recognition of others by biological motion: viewpoint-dependent effects. Perception 35, 911–920.

81. Little, J., and Boyd, J. (1998). Recognizing people by their gait: The shape of motion.

82. Gafurov, D., and Snekkenes, E. (2009). Gait Recognition Using Wearable Motion Recording Sensors. EURASIP J. Adv. Signal Process. 2009, 415817.

83. Makihara, Y., Nixon, M.S., and Yagi, Y. (2020). Gait Recognition: Databases, Representations, and Applications. In Computer Vision: A Reference Guide (Springer International Publishing), pp. 1–13.

84. Ernst, M., and Fudge, J.L. (2009). A developmental neurobiological model of motivated behavior: anatomy, connectivity and ontogeny of the triadic nodes. Neurosci. Biobehav. Rev. 33, 367–382.

85. Li, Y., and Dulac, C. (2018). Neural coding of sex-specific social information in the mouse brain. Curr. Opin. Neurobiol. 53, 120–130.

86. Ferguson, J.N., Aldag, J.M., Insel, T.R., and Young, L.J. (2001). Oxytocin in the medial amygdala is essential for social recognition in the mouse. J. Neurosci. 21, 8278–8285.

87. Young, L.J. (2002). The neurobiology of social recognition, approach, and avoidance. Biol. Psychiatry 51, 18–26.

88. Ko, J. (2017). Neuroanatomical Substrates of Rodent Social Behavior: The Medial Prefrontal Cortex and Its Projection Patterns. Front. Neural Circuits 11, 41.

89. Newman, S.W. (1999). The medial extended amygdala in male reproductive behavior. A node in the mammalian social behavior network. Ann. N. Y. Acad. Sci. 877, 242–257.

90. Raam, T., and Hong, W. (2021). Organization of neural circuits underlying social behavior: A consideration of the medial amygdala. Curr. Opin. Neurobiol. 68, 124–136.

91. Rasia-Filho, A.A., Londero, R.G., and Achaval, M. (2000). Functional activities of the amygdala: an overview. J. Psychiatry Neurosci. 25, 14–23.

92. Martersteck, E.M., Hirokawa, K.E., Evarts, M., Bernard, A., Duan, X., Li, Y., Ng, L., Oh, S.W., Ouellette, B., Royall, J.J., et al. (2017). Diverse Central Projection Patterns of Retinal Ganglion Cells. Preprint, 10.1016/j.celrep.2017.01.075 10.1016/j.celrep.2017.01.075.

93. Elliott, A.S., Weiss, M.L., and Nunez, A.A. (1995). Direct retinal communication with the peri-amygdaloid area. Neuroreport 6, 806–808.

94. Morin, L.P., and Studholme, K.M. (2014). Retinofugal projections in the mouse. J. Comp. Neurol. 522, 3733–3753.

95. Delwig, A., Larsen, D.D., Yasumura, D., Yang, C.F., Shah, N.M., and Copenhagen, D.R. (2016). Retinofugal Projections from Melanopsin-Expressing Retinal Ganglion Cells Revealed by Intraocular Injections of Cre-Dependent Virus. PLoS One 11, e0149501.

96. Cádiz-Moretti, B., Otero-García, M., Martínez-García, F., and Lanuza, E. (2016). Afferent projections to the different medial amygdala subdivisions: a retrograde tracing study in the mouse. Brain Struct. Funct. 221, 1033–1065.

97. Johnson, R.F., Morin, L.P., and Moore, R.Y. (1988). Retinohypothalamic projections in the hamster and rat demonstrated using cholera toxin. Brain Res. 462, 301–312.

98. Hattar, S., Kumar, M., Park, A., Tong, P., Tung, J., Yau, K.-W., and Berson, D.M. (2006). Central projections of melanopsin-expressing retinal ganglion cells in the mouse. J. Comp. Neurol. 497, 326–349.

99. Martersteck, E.M., Hirokawa, K.E., Evarts, M., Bernard, A., Duan, X., Li, Y., Ng, L., Oh, S.W., Ouellette, B., Royall, J.J., et al. (2017). Diverse Central Projection Patterns of Retinal Ganglion Cells. Cell Rep. 18, 2058–2072.

100. Levine, J.D., Weiss, M.L., Rosenwasser, A.M., and Miselis, R.R. (1991). Retinohypothalamic tract in the female albino rat: a study using horseradish peroxidase conjugated to cholera toxin. J. Comp. Neurol. 306, 344–360.

101. Pu, M. (1999). Dendritic morphology of cat retinal ganglion cells projecting to suprachiasmatic nucleus. J. Comp. Neurol. 414, 267–274.

102. Mick, G., Cooper, H., and Magnin, M. (1993). Retinal projection to the olfactory tubercle and basal telencephalon in primates. J. Comp. Neurol. 327, 205–219.

103. McCarthy, M.M., and Vom Saal, F.S. (1986). Inhibition of infanticide after mating by wild male house mice. Physiol. Behav. 36, 203–209.

104. Kondo, Y., and Sachs, B.D. (2002). Disparate effects of small medial amygdala lesions on noncontact erection, copulation, and partner preference. Physiol. Behav. 76, 443–447.

105. Domínguez-Borràs, J., Moyne, M., Saj, A., Guex, R., and Vuilleumier, P. (2020). Impaired emotional biases in visual attention after bilateral amygdala lesion. Neuropsychologia 137, 107292.

106. Young, L.J., and Zhang, Q.I. (2021). On the Origins of Diversity in Social Behavior. 動物心理学研究 71, 45–61.

107. O’Connell, L.A., and Hofmann, H.A. (2011). Genes, hormones, and circuits: an integrative approach to study the evolution of social behavior. Front. Neuroendocrinol. 32, 320–335.

108. Zhou, T., Sandi, C., and Hu, H. (2018). Advances in understanding neural mechanisms of social dominance. Curr. Opin. Neurobiol. 49, 99–107.

109. Bordes, J., Miranda, L., Müller-Myhsok, B., and Schmidt, M.V. (2023). Advancing social behavioral neuroscience by integrating ethology and comparative psychology methods through machine learning. Neurosci. Biobehav. Rev. 151, 105243.

110. Mack, N.R., Bouras, N.N., and Gao, W.-J. (2024). Prefrontal regulation of social behavior and related deficits: insights from rodent studies. Biol. Psychiatry. 10.1016/j.biopsych.2024.03.008.

111. Kim, Y., Venkataraju, K.U., Pradhan, K., Mende, C., Taranda, J., Turaga, S.C., Arganda-Carreras, I., Ng, L., Hawrylycz, M.J., Rockland, K.S., et al. (2015). Mapping social behavior-induced brain activation at cellular resolution in the mouse. Cell Rep. 10, 292–305.

112. Anderson, D.J. (2016). Circuit modules linking internal states and social behaviour in flies and mice. Nat. Rev. Neurosci. 17, 692–704.

113. Zilkha, N., Sofer, Y., Beny, Y., and Kimchi, T. (2016). From classic ethology to modern neuroethology: overcoming the three biases in social behavior research. Curr. Opin. Neurobiol. 38, 96–108.

114. Kuti, O.J., and Page, D.T. (2011). Assessment of Social Approach Behavior in Mice. In Mood and Anxiety Related Phenotypes in Mice: Characterization Using Behavioral Tests, Volume II, T. D. Gould, ed. (Humana Press), pp. 83–95.

115. Powell, S.B., Geyer, M.A., Gallagher, D., and Paulus, M.P. (2004). The balance between approach and avoidance behaviors in a novel object exploration paradigm in mice. Behav. Brain Res. 152, 341–349.

116. Bechtholt, A.J., Gremel, C.M., and Cunningham, C.L. (2004). Handling blocks expression of conditioned place aversion but not conditioned place preference produced by ethanol in mice. Pharmacol. Biochem. Behav. 79, 739–744.

117. Fulenwider, H.D., Caruso, M.A., and Ryabinin, A.E. (2022). Manifestations of domination: Assessments of social dominance in rodents. Genes Brain Behav. 21, e12731.

118. Goyens, J., and Noirot, E. (1975). Effects of cohabitation with females on aggressive behavior between male mice. Dev. Psychobiol. 8, 79–84.

119. Brain, P.F., Benton, D., and Bolton, J.C. (1978). Comparison of agonistic behavior in individually-housed male mice with those cohabiting with females. Aggress. Behav. 4, 201–206.

120. Wegner, K.A., Abler, L.L., Oakes, S.R., Mehta, G.S., Ritter, K.E., Hill, W.G., Zwaans, B.M., Lamb, L.E., Wang, Z., Bjorling, D.E., et al. (2018). Void spot assay procedural optimization and software for rapid and objective quantification of rodent voiding function, including overlapping urine spots. Am. J. Physiol. Renal Physiol. 315, F1067–F1080.

121. Schindelin, J., Arganda-Carreras, I., Frise, E., Kaynig, V., Longair, M., Pietzsch, T., Preibisch, S., Rueden, C., Saalfeld, S., Schmid, B., et al. (2012). Fiji: an open-source platform for biological-image analysis. Nat. Methods 9, 676–682.

122. Coffey, K.R., Marx, R.E., and Neumaier, J.F. (2019). DeepSqueak: a deep learning-based system for detection and analysis of ultrasonic vocalizations. Neuropsychopharmacology 44, 859–868.

123. Friard, O., and Gamba, M. (2016). BORIS: a free, versatile open-source event-logging software for video/audio coding and live observations. Methods Ecol. Evol. 7, 1325–1330.

