## Supplemental information for "Visual Identification of Conspecifics Shapes Social Behavior in Mice"

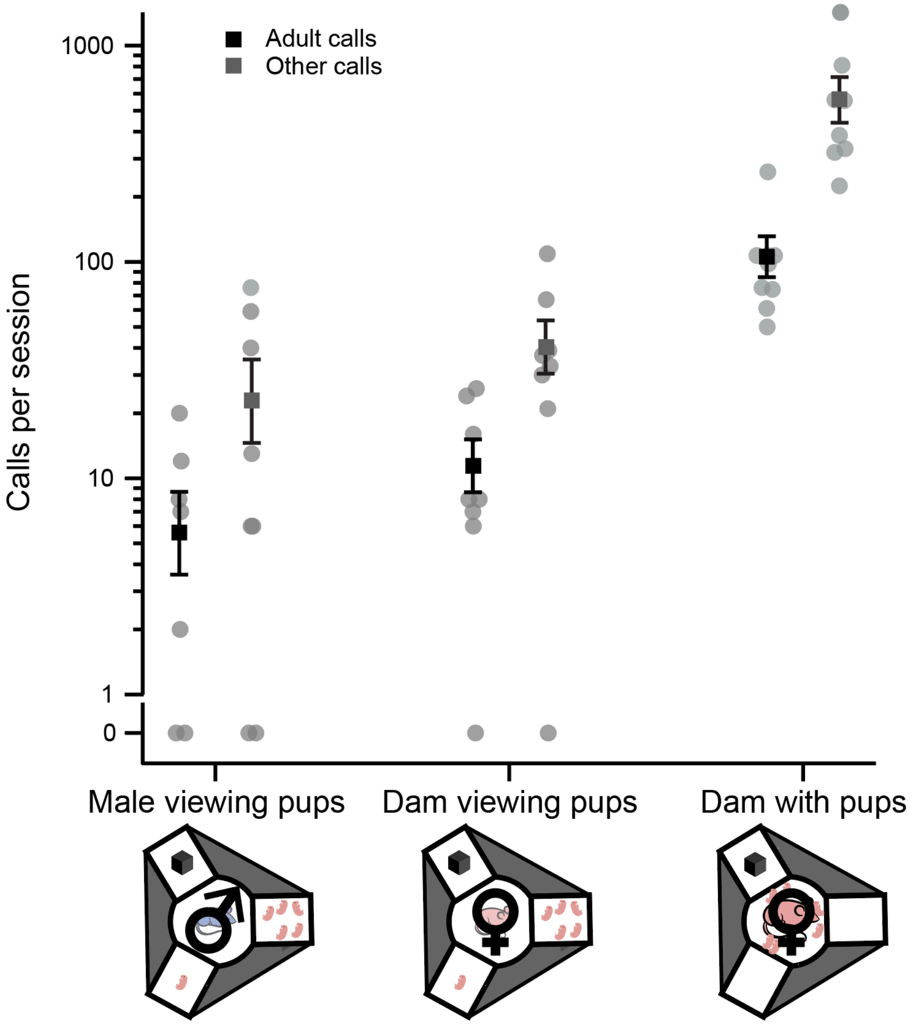


**Figure S1 Total USV comparisons between the conditions.** Call numbers in conditions with a lone male viewing pups and a novel object, compared to a dam viewing pups and a novel object, and a dam and pups viewing a novel object. The same number of pups in the stimulus chamber were in the arena (n = 6). The difference between the dam and pups viewing the novel object compared to the dam viewing pups/novel object was $p = 0.0043$ between total calls of dams vs dams/pups.

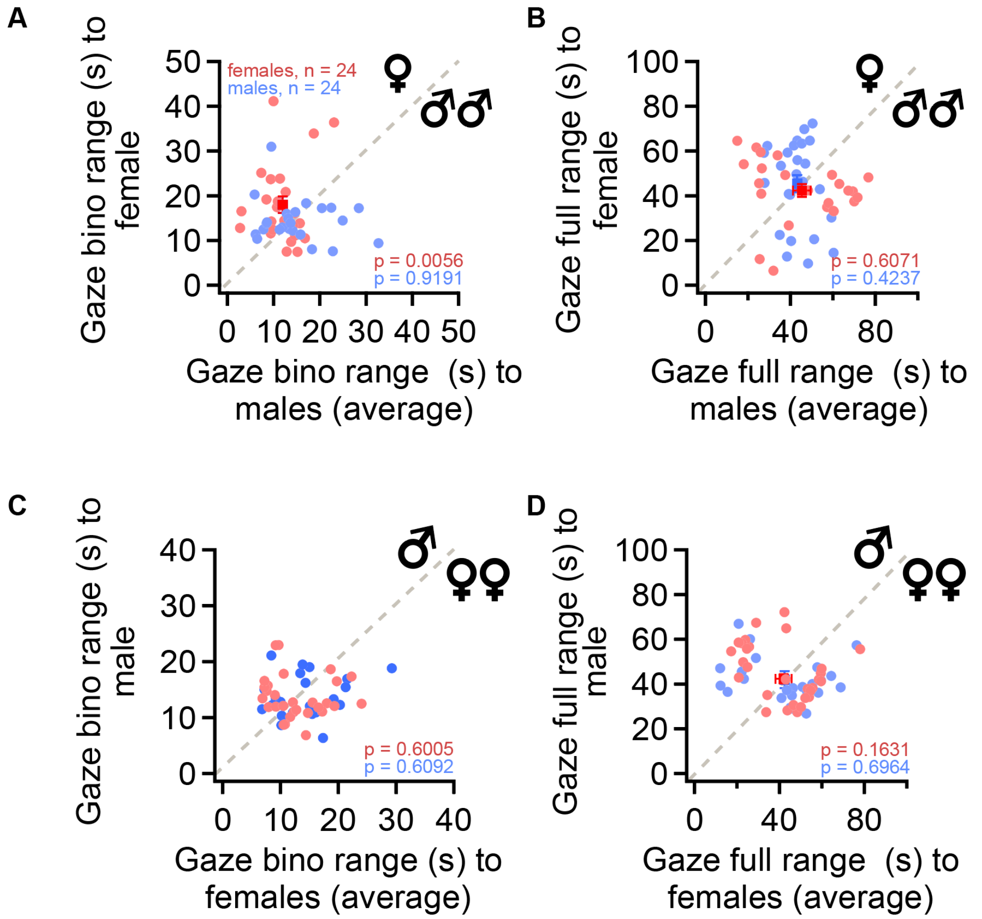


**Figure S2 The binocular gaze range and full gaze range of mice during sex discrimination. (A)** The number of seconds each mouse kept a stimulus mouse within the binocular gaze range of their sight for the one female and two male condition. **(B)** For the same experiment, the number of seconds each mouse spent with the stimulus mouse within the line of their full gaze range. **(C-D)** Same parameters explored but for the one male and two female stimuli condition.

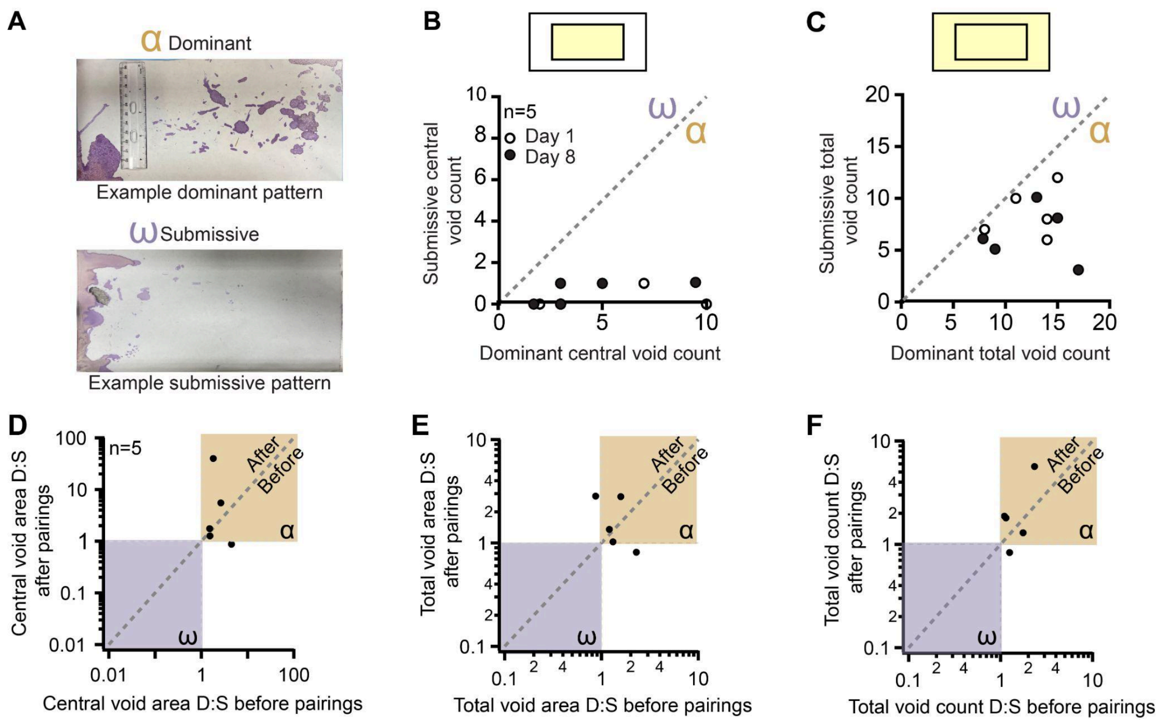


**Figure S3 Metrics for monitoring the social dominance within a cage through void patterns. (A)** Upper panel: example dominant voiding pattern on the bottom of a cage. Lower panel: example submissive voiding pattern. **(B)** Void count in the center of the arena between the dominant and submissive of each hierarchy, open marker for Day 1 of experiment and closed for Day 8. **(C)** Total count of urinations in the arena between the dominant and submissive mice. **(D-F)** Comparison between the Dom:Sub ratio for voids in the center of the arena **(D)**, number of void+ pixels in all of arena **(E)**, number of voids in total **(F)** before and after dominance manipulations. Purple arena indicates a submissive leading result, orange arena for dominant leaning.

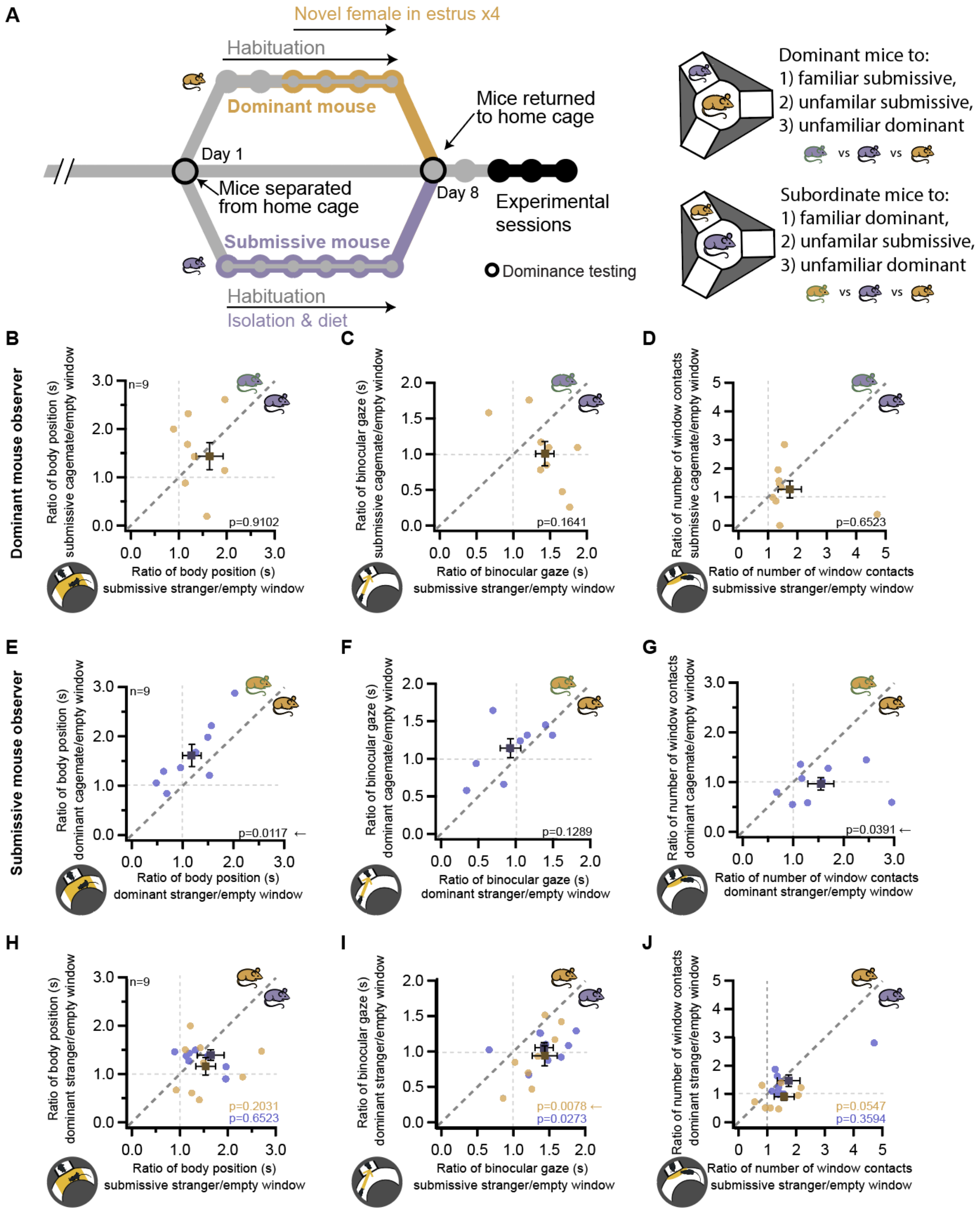


**Figure S4 Mouse behavior depends on the visual identification of the social status of a stranger mouse.**

**(A)** Left panel: Mouse cohort dominance experiment setup. One dominant mouse is separated from the cage and manipulated to maintain their dominance in the cohort. Submissive mice are isolated and put on a dietary restriction. Right panel: Dominant mice (orange, n = 9) view in separate experiments a familiar submissive cagemate (green and purple), an unfamiliar submissive mouse (purple), or an unfamiliar dominant mouse (orange). Submissive mice (purple, n = 9) view their dominant cagemate, a dominant stranger, or a submissive stranger. **(B-D)** The proportion of time near, gaze facing, or contacts made towards a submissive cagemate over the adjoining empty windows compared to the ratio of those same metrics in sessions with a submissive stranger (body position $p = 0.9102$, gaze $p = 0.0273$, and window contacts $p = 0.6523$). **(E-G)** For submissive mice, the proportion of time spent nearby, facing, or contacting their dominant cagemate's window vs the equivalent for dominant stranger conditions. **(H-J)** The proportion of time nearby, gazing towards, or contacting the window of a dominant stranger versus a submissive stranger (body position $p = 0.2031$, head gaze $p = 0.0078$, window contacts $p = 0.0547$). The proportion of time nearby, gazing towards, or contacting the window of a dominant stranger versus a submissive stranger (body position $p = 0.6523$, head gaze $p = 0.0273$, window contacts $p = 0.3594$).

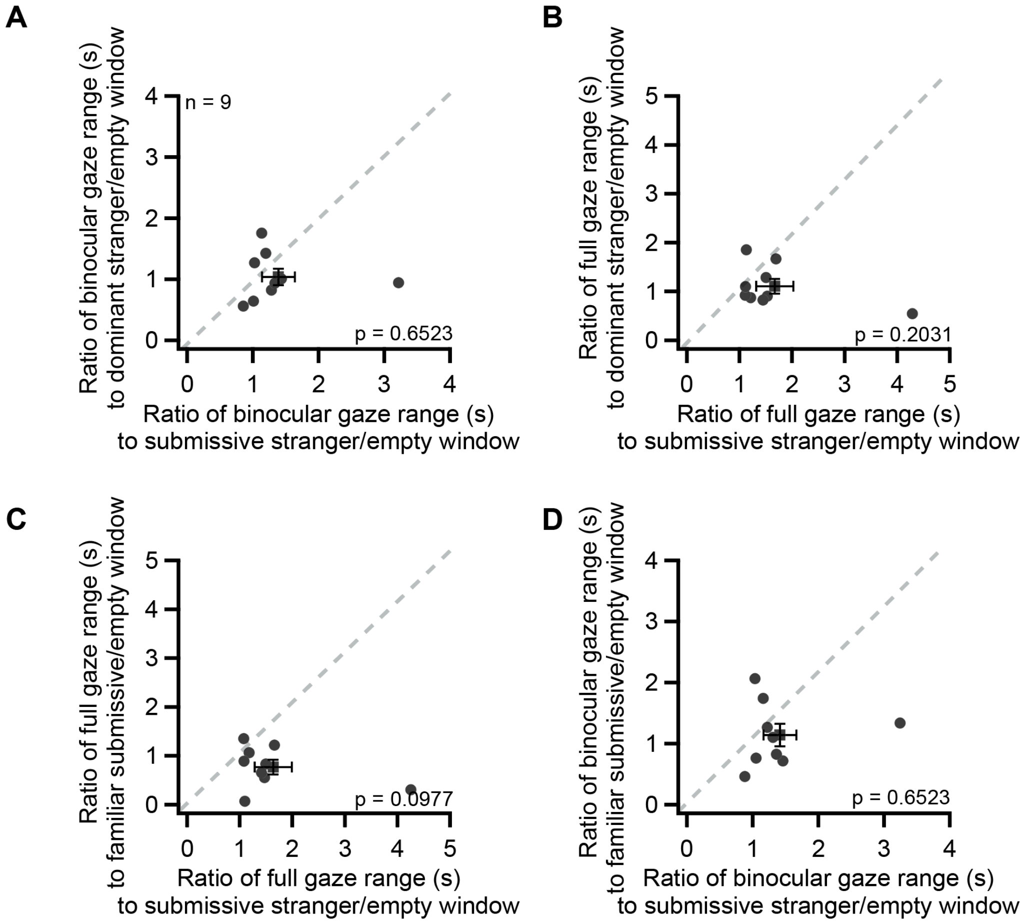


**Figure S5 The binocular gaze range and full gaze range of dominant mice during the familiarity and status experiments. (A)** The number of seconds each mouse kept a stimulus mouse within the binocular gaze range of their sight for the dominant and submissive strangers conditions. **(B)** For the same experiment, the number of seconds each mouse spent with the stimulus mouse within the line of their full gaze range. **(C-D)** Same parameters explored but for the familiar submissive compared to submissive stranger conditions.

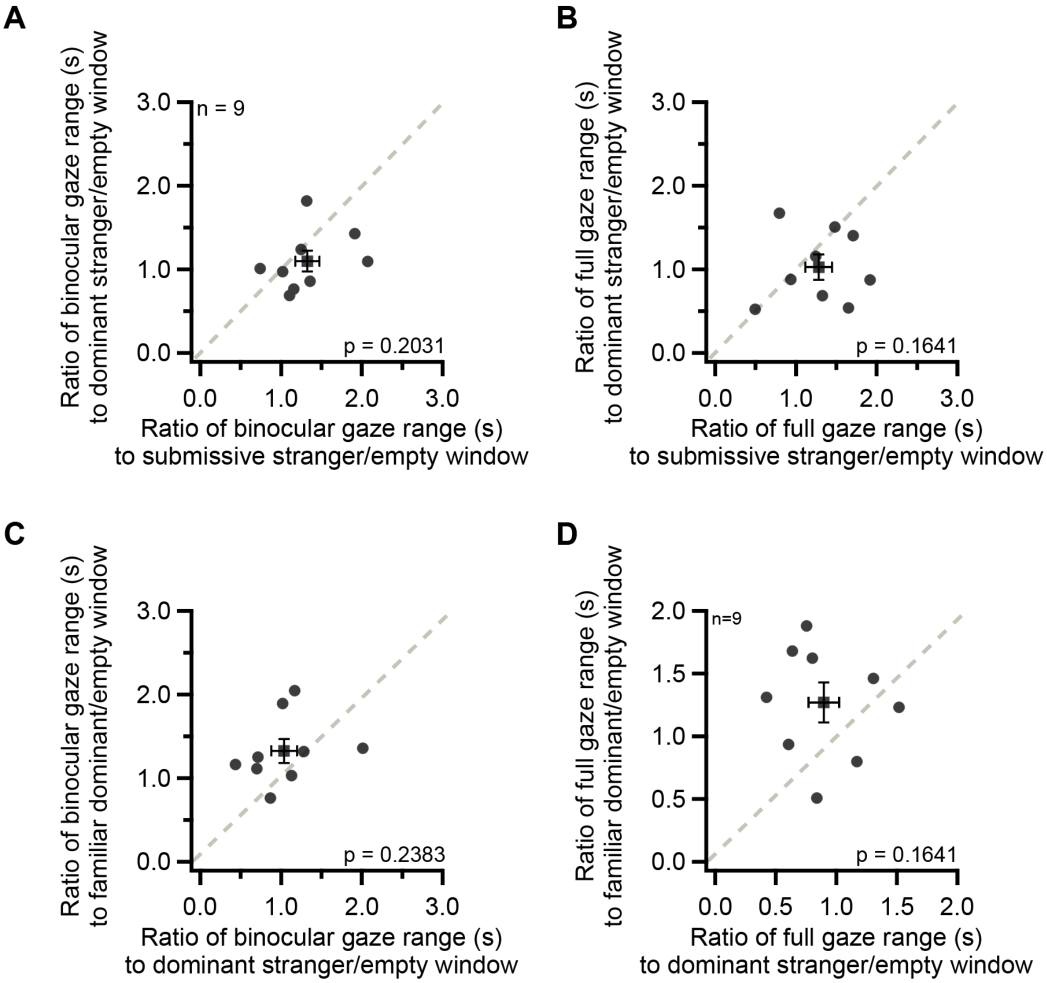


**Figure S6 The binocular gaze range and full gaze range of submissive mice during the familiarity and status experiments. (A)** The number of seconds each mouse kept a stimulus mouse within the binocular gaze range of their sight for the dominant and submissive strangers conditions. **(B)** For the same experiment, the number of seconds each mouse spent with the stimulus mouse within the line of their full gaze range. **(C-D)** Same parameters explored but for the familiar dominant compared to dominant stranger conditions.
